# Indirect Haptic Disturbances Enhance Motor Variability, with Divergent Effects on Skill Transfer

**DOI:** 10.1101/2024.02.29.582677

**Authors:** Wouter Arink, Katherine L. Poggensee, Niek Beckers, David A. Abbink, Laura Marchal-Crespo

## Abstract

Research on motor learning has found evidence that learning rate is positively correlated with the learner’s motor variability. However, it is still unclear how to robotically promote that variability without compromising the learner’s sense of agency and motivation, which are crucial for motor learning. We propose a novel method to enhance motor variability during learning of a dynamic task by applying pseudorandom perturbing forces to the internal degree of freedom of the dynamic system rather than directly applying the forces to the learner’s limb. Twenty healthy participants practiced swinging a virtual pendulum to hit oncoming targets, either with the novel method or without disturbances, to evaluate the effect of the method on motor learning, skill transfer, motivation, and agency. We evaluated skill transfer using two tasks, changing either the target locations or the task dynamics by shortening the pendulum rod. The indirect haptic disturbance method successfully increased participants’ motor variability during training compared to training without disturbance. Although we did not observe group-level differences in learning, we observed divergent effects on skill generalization. The indirect haptic disturbances seemed to promote skill transfer to the altered task dynamics but limited transfer in the task with altered target positions. Motivation was not affected by the haptic disturbances, but future work is needed to determine if indirect haptic noise negatively impacts sense of agency. Increasing motor variability by indirect haptic disturbance is promising for enhancing skill transfer in tasks that incorporate complex dynamics. However, more research is needed to make indirect haptic disturbance a valuable tool for real-life motor learning situations.

## I. Introduction

Motor learning is a process we all encounter throughout our lives, from learning how to walk as a baby to practicing music, and provides a framework to study and improve neurorehabilitation. Motor variability, the variability in movements made when performing a motor task, plays an important role in motor learning. While early theories considered motor variability to be a reflection of undesired noise from the nervous system [1]–[3], current understanding suggests that the brain actively regulates variability to facilitate learning [4]. Yet, there is currently no consensus on how induced variability affects motor learning.

Haptic forces, applied as error amplification or random disturbances, have been used to promote learning by increasing motor variability [5]. However, direct forces applied dur-ing task execution might decrease the participants’ feelings of competence [6], [7] and motivation [8], and therefore, potentially hamper motor learning [9]. Indirect haptic forces might offer a solution to enhance motor variability without hampering these crucial psychological factors. By applying forces to an internal degree of freedom of the system to be learned, participants may only indirectly perceive those forces, thus mitigating the negative effects on participants’ sense of agency [10] and motivation.

A crucial element in motor learning, that is often not thoughtfully evaluated, is the transfer of the learned skills to similar, albeit different, tasks. The primary goal in real life is usually a broader scope of tasks, varying in task dynamics (e.g., different golf clubs) or targets (e.g., different distances to the hole), rather than improvement in only the—usually non-ecologically valid—trained task in robot-aided training. Yet, the effect of increased variability during robotic training on skill transfer has been hardly evaluated [5].

In this study, we investigated the effect of applying indirect haptic perturbations on the internal degree of freedom of a pendulum on motor learning. The task consisted of controlling a virtual pendulum to hit oncoming targets. We analyzed the hand movement variability and the variability of the internal degree of freedom, hypothesizing that this haptic strategy would drive participants to increase motor variability during training compared to unperturbed participants. We examined if this increased motor variability led to learning benefits in the trained task and evaluated the transfer of the gained skills with two different tasks: one with a slightly different dynamic system, and a second one that required a different movement planning. We also measured participants’ sense of agency and motivation, expecting no effect from the indirect haptic disturbance.

## II. Methods

### A. Participants and Experimental Setup

Twenty participants (nine female, two left-handed, age range of 18-26) participated in the study approved by the TU Delft Ethics Committee.

Participants sat in front of a monitor with a haptic device (Delta.3, Force Dimension, Switzerland) positioned on the side of their dominant hand while resting their elbow on the table (Fig. 1A). The robot motion control and task visualization, implemented in C++ and Unity (Unity Technologies, US) respectively, were adapted from Özen et al. [11].

**Fig. 1:**
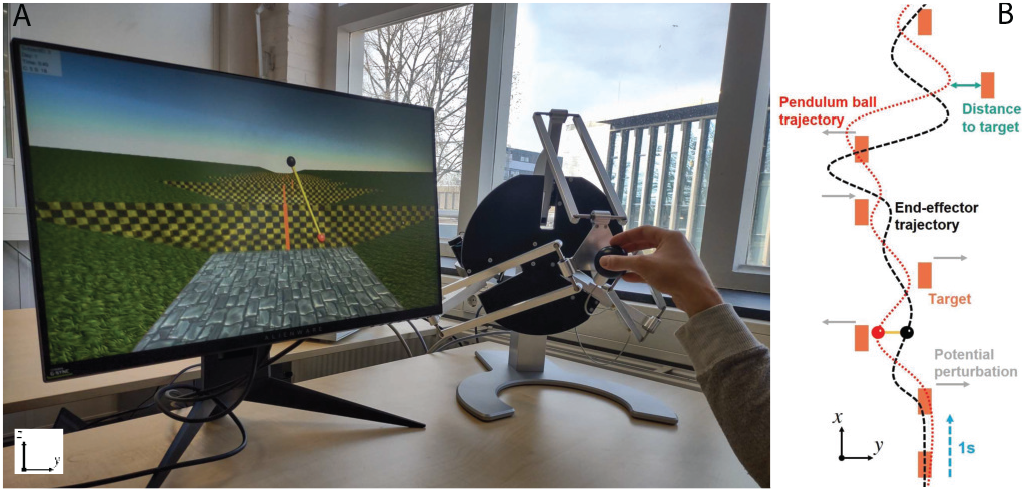
A: Experiment setup. Participants sat in front of the monitor next to the haptic Delta.3 device, on the side of their dominant hand. The pendulum ball (red dot) was controlled by moving the end-effector of the robot, which is represented by the black dot on the screen (pendulum pivoting point). With respect to the screen, the *z* axis represents vertical movement, and horizontal movement occurs along the *y* axis. B: Schematic top view (*z*plane) of a trial with exemplary trajectories of the ball (red dotted) and end effector (black dashed). The gray arrows indicate the potential perturbation locations. Note that at most three perturbations were applied during a trial and never after two consecutive targets.

### B. The Pendulum Game

The task consisted of moving a virtual pendulum’s pivot point shown on the screen by moving the end-effector of the haptic device (Fig. 1A), with a one-to-one mapping, to hit incoming targets with the pendulum ball. By moving the pendulum pivot point, participants controlled the pendulum ball position (red ball in Fig. 1A&B) following the equation of motion of a single pendulum (see [11]). Participants were instructed to hit oncoming vertical targets with different horizontal positions with the pendulum ball as accurately as possible. One trial included eight targets separated by 1 s (Fig. 1B). A period of 5 s without targets followed each trial after which the next trial began to give participants time to return to the starting position. The workspace of the end-effector was constrained to move only in the vertical *x*-plane. Importantly, the haptic device rendered the forces from the dynamics of the pendulum back to the end-effector [12].

A score was given after each target, based on the distance along the *y*-direction between the pendulum ball and the center of the target (Fig. 1B, [11]) with a 0.5 point deduction from 100 for every mm deviation from the target. Scores were shown for 0.5 s, in green for positive scores and red for zero, with an average trial score displayed at the end of the trial.

#### 1) Main and transfer tasks

Participants practiced three target-hitting tasks across two days. During the main task, targets were positioned to increase task difficulty throughout the trial, starting with two targets in the center of the workspace, followed by three targets varying by 30 mm from left to right (*y*-direction) and targets 6–8 placed to exploit the considerable swing and distance covered by the previous targets. To make the task non-repetitive and thus engaging, in half of the trials the target locations were mirrored about the *x*-axis. The order of mirrored / non-mirrored trials was pseudo-randomized, with the same order for every participant.

Two different transfer tasks were also implemented. The first transfer task consisted of hitting targets with the same pendulum that appeared in different locations than those in the main task. The difficulty of this transfer task was increased with respect to the main task by increasing the horizontal distance between the targets and the centerline by 5 mm. For the second transfer task, the positioning of the targets remained the same as in the main task, but the pendulum dynamics were altered by decreasing the length of the pendulum by 30 %. Decreasing the rod length increases the natural frequency of the pendulum, making the pendulum swing faster. The pendulum length was not visually changed on the screen, so the altered dynamics could only be perceived by changes in how the pendulum behaved and felt through the haptic rendering.

### C. Experimental Protocol

The experiment took place across two days, separated by 1-3 days (Fig. 2). The first day started with a familiarization period, during which participants familiarized themselves with the haptic device and pendulum dynamics with four trials from the main task, with the even targets mirrored. This familiarization was preceded by 40 s of moving the pendulum without targets.

**Fig. 2:**
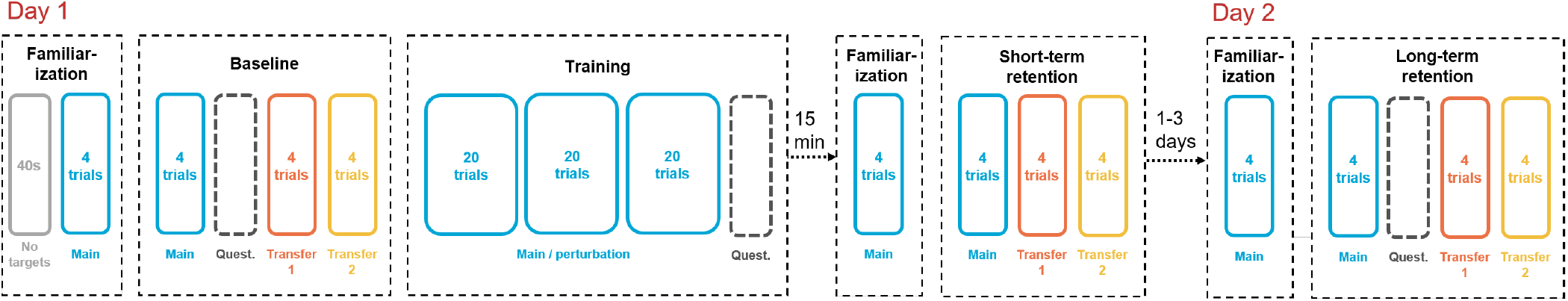
Study Protocol. Participants were randomly allocated to either the control or the experimental group that received perturbation forces during training.

Performance metrics and questionnaires (see section II- D) were taken during the testing periods, i.e., baseline (BL), short-term retention (STR) —performed 15 min after training—-, and long-term retention tests (LTR)—performed on the second day. Both retention tests were preceded by a shortened (re)familiarization period. Each one of these testing periods included four trials for each task, i.e., the main task and the two transfer tasks, always in the same order.

Between BL and STR, participants trained the main task with three blocks of twenty trials each, separated by optional rest periods. Each block was identical in terms of the order of the mirrored trials. Participants were randomly allocated to two training groups (Control and Experimental, ten per group, five females, and one left-handed in the control group). The control group (C) trained without any perturbation from the robot (only with the haptic rendering of the pendulum dynamics), and the experimental group (P) experienced perturbation forces applied on the pendulum ball tangentially to its movement during training on top of the haptic rendering. Per trial, two or three perturbation forces were pseudo-randomly applied directly to the pendulum ball in the opposite direction of the location of the next target. Several components of the perturbations were varied from trial to trial to prevent participants from anticipating the perturbations, such that every participant in this group trained with the same settings in the same order. The onset of the perturbation was pseudo-randomly selected from 20- 80 ms after crossing a target and lasted for 50 ms to give participants time to compensate for the perturbation before crossing the next target. The magnitude of the perturbation forces also varied from 0.8-1.5 N. The mean magnitude and onset time of the perturbation forces decreased over the trial as the difficulty posed by the position of the targets increased.

Trials 3, 7, 13, and 17 in each block were catch trials (i.e., no perturbations were applied) to evaluate the effect of the perturbation forces on participants’ motor variability compared to unperturbed trials.

### D. Data Processing

Several metrics were selected to evaluate motor variability, motor learning, and sense of agency and motivation.

#### 1) Motor variability

We selected three different measures of variability. To see if the perturbations affected the variability of the pendulum’s internal degree of freedom, we analyzed participants’ swing variability by taking the standard deviation of the pendulum angle for each trial. To quantify the participants’ end-effector workspace exploration, the end-effector variability was calculated as the standard deviation of the horizontal end-effector (i.e., hand) position. Since the task already consists of inherent end-effector variability, as horizontal positions of the targets vary, we also quantified how much participants explored from trial to trial over the entire training by evaluating the path variability, defined as the standard deviation of the end-effector horizontal position around the participant’s mean path. The participant’s mean path was computed by taking the mean *y*-position of every time instant of a trial over all sixty trials of the training block.

#### 2) Motor learning

We used three different performance metrics to evaluate motor learning. First, the trial score was the average score of the eight targets within a trial. Participants’ score variability within a trial was calculated to evaluate how consistent they were at hitting all the targets and is expected to negatively correlate with skill level [11], [13].

We further evaluated if participants learned to control the pendulum dynamics by quantifying the deviation from the pendulum natural frequency, which was previously found to correlate with performance [11]. The target positions do not follow a periodic order, so participants needed to adjust the swing frequency away from its natural frequency. We computed the power spectral density (PSD) of the swing angle and the percentage of power at the pendulum natural frequency (*ω*_*n*_=0.57 Hz) for each trial (similar to [11] and [14]). As the duration of a trial was 8 s, the frequency resolution was 0.125 Hz. We therefore selected the power at frequencies 0.5 and 0.625 Hz as the power at the natural frequency. For the second transfer task with different pendulum length, the natural frequency was *ω*_*n*_=0.69 Hz, so we selected the power at 0.625 and 0.75 Hz as the power at natural frequency.

#### 3) Psychological Factors

Participants filled out two questionnaires after BL, training, and LTR (Fig. 2). To determine how the perturbations affected participants’ feelings of control over the pendulum’s movements, we assessed their sense of agency with three adapted questions from the questionnaire used by Piryankova et al. [10]. Each question was scored from -3 (strongly disagree) to 3 (strongly agree). Participants’ motivation was evaluated using questions from the Intrinsic Motivation Inventory (IMI) Questionnaire [15]. Participants answered 12 questions on a scale from 1 (not at all true) to 7 (very true) related to their: *Interest/Enjoyment, Perceived Competence, Effort/Importance*, and *Pressure/Tension*.

### E. Statistical Analysis

We performed a regression analysis on linear mixed effects (lme) models to evaluate the effect of the perturbations on motor variability and learning. The pymer4 package from *R* in Python was used for the statistical analysis. The models were fitted using a maximum likelihood estimation. The confidence intervals (CI) of the estimated coefficients were computed using bootstrapping (ten thousand iterations) [16].

The following variables were included in the analysis. *Group* (*Control, Experimental*) and *Catch* trial (*Y es* = 1, *No* = 0) are categorical variables with two levels each. Note that *Catch* = 1 for all trials in the Control group. *Time* is a categorical variable with three levels: BL, STR, and LTR. *Participant* was included as a random factor. To evaluate if the perturbing forces increased motor variability during training, we modeled the three motor variability metrics as a function of the interaction between *Group* and *Catch* with *Participant* as a random factor. All trials from the three training blocks were included, except for those with a path variability outside three standard deviations from a participant’s mean path.

To evaluate the effect of training on motor learning, we modeled each of the three performance metrics as a function of the interaction between *Group* and *Time*, with *Time*|*Participant* included as a random slope to account for individual differences in learning ability. For every block, all four trials were used for the analysis. This model was also used with the motor variability metrics as the dependent variable to check for group-level differences at Baseline.

To evaluate the effect of the perturbations on the psychological factors, we modeled the average scores per subscale from the IMI and agency questionnaires as a function of the interaction between *Group* and *Time* and the random factor *Participant*.

Finally, we computed the repeated-measures correlation [17] to evaluate if there was a relationship between the score and the deviation from the pendulum natural frequency.

## III. Results

We found no significant differences between groups for any of the motor variability or performance metrics in the main task BL block. We found a significant correlation between the score and the power at natural frequency, *r* = −0.21 (95 % CI [−0.25, −0.16]).

### A. Motor Variability During Training

For all three motor variability metrics, there was a significant effect of group, indicating that the experimental group P showed higher variability during training than the control group (Table I). For the pendulum swing variability and horizontal end-effector variability, there was also a significant interaction between *Group* and *Catch*, indicating that the perturbations significantly increased these two types of variability in perturbation trials compared to catch trials.

**TABLE 1:**
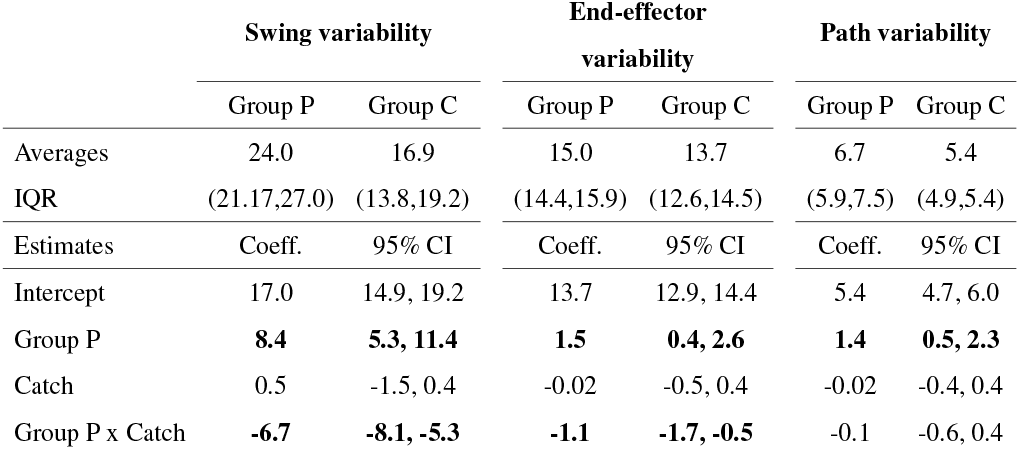
Motor variability during training: averages (with interquartile ranges) and results from the linear regression analysis. Significant effects are printed in bold. P: Experimental group, C: Control group.

### B. Motor Learning

In general, participants increased their performance significantly from BL to STR and LTR for all three performance metrics, although the reduction in power around the pendulum natural frequency from BL to STR was only trending towards significance (Fig. 3, Table II). There was, however, no significant interaction between *Time* and *Group* for both STR and LTR for any of the performance metrics, suggesting no learning differences between groups in the main task. We found similar significant improvements in performance in the two transfer tasks. However, in both transfer tasks, there was a significant interaction between *Group* and *Time* during the retention tests. For transfer task 1 (altered target positions), the control group decreased the power around the pendulum’s natural frequency significantly more than the experimental group at STR and LTR, while in transfer task 2 (reduced rod length), the opposite trend was observed. For the latter, this interaction was only observed from BL to STR.

**Fig. 3:**
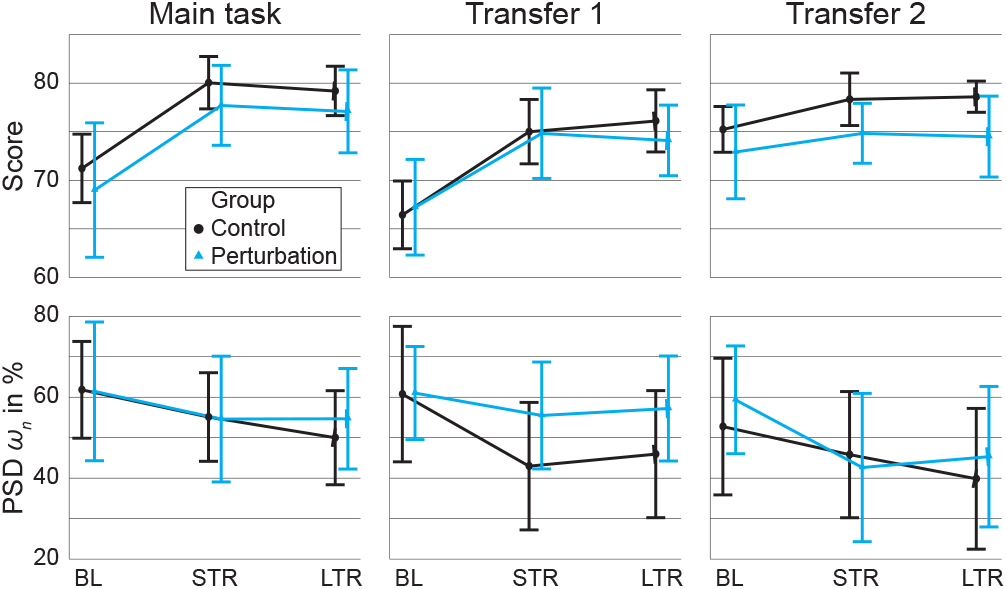
Average score (top) and average power around the pendulum natural frequency (bottom) of the two groups in baseline (BL), short-term (STR) and long-term retention (LTR) of the main task, transfer task with altered target positions (Transfer 1) and transfer task with reduced pendulum length (Transfer 2). The error bars represent one standard deviation from the mean.

**TABLE 2:**
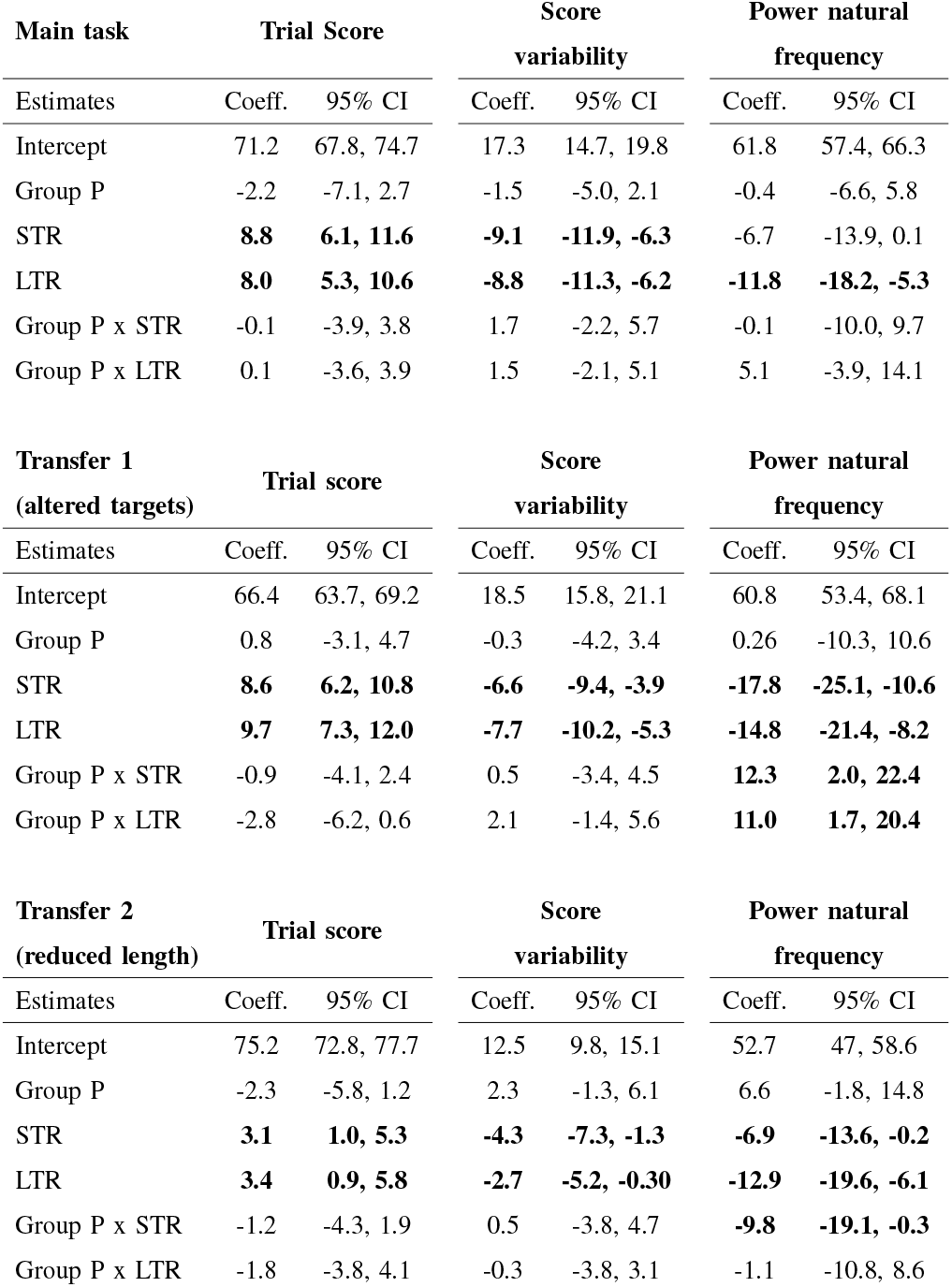
Motor learning evaluation. Results from the linear regression analysis. Significant effects are printed in bold. P: Experimental group, STR: Short-term Retention, LTR: Long-term Retention.

### C. Psychological Factors

The statistical analysis did not reveal any significant differences between the groups in participants’ reported sense of agency or motivation subscales.

Participants’ perceived sense of agency significantly increased from BL to LTR (0.6, 95 %CI [0.03,1.2]). Furthermore, the interaction between *Group* and *Time* was trending towards significance from BL to training (−0.7, 95 %CI [−1.5, 0.1]), which indicated that participants’ sense of agency may have decreased due to the perturbations.

No significant interaction effects between *Group* and *Time* were found in any of the motivation subscales. There was also no significant effect of *Time* for the subscales *Interest/Enjoyment* and *Effort/Importance*, which remained high through the experiment (5.5 and 5.4 out of 7 at BL, respectively). Participants’ *Perceived Competence*, on the other hand, significantly increased from BL to training and LTR (1.8, 95 %CI [1.1, 2.5] and 1.5, 95 %CI [0.8, 2.2], respectively).

## IV. Discussion

We studied the effect on motor learning of a new haptic method to increase participants’ motor variability during training by applying pseudo-random perturbation forces to the internal degree of freedom of the system to be controlled (indirect haptic disturbance). These perturbations had the expected effect of increasing motor variability. While the perturbations did not lead to an improved score compared to the control group, we did observe differences between groups in participants’ ability to swing the pendulum away from the natural frequency in the transfer tasks, which is related to better performance. This suggests a benefit of training with indirect haptic perturbations for transfer of learning to altered task dynamics, but a disadvantage for transfer of learning to altered target positions.

### A. Indirect haptic perturbation increases motor variability

The perturbing forces directly on the pendulum’s ball had the desired effect of increasing motor variability in all three metrics. Therefore, applying haptic disturbances to the internal degree of freedom of a dynamic system is a valid method for increasing motor variability, rather than applying the disturbance directly to the participants’ limbs [6].

The path variability metric indicates the deviation from the participant’s main path, so it is unsurprising that the interaction between *Group* and *Catch* was not significant. The perturbations are expected to considerably influence the mean path. Therefore, the catch trials would also differ more from a participant’s mean path for participants training with perturbations compared to participants in the control group.

We focused only on the horizontal direction in the analysis of the variability of the end-effector and its path. It is possible that the perturbations also caused differences in motor variability in the vertical *z*-direction, although this information could be less important as target positions and corresponding scores only varied in the horizontal direction.

### B. Indirect haptic disturbance does not affect motor learning of the trained task

We observed no differences between the training groups in learning the trained task, despite the enhanced variability exhibited by the experimental group during training. This is in line with previous literature that did not find advantages of increased variability with direct perturbations on motor learning [6], [7]. While these previous studies hypothesized that the lack of benefit of haptic disturbance might be due to a reduction in participants’ motivation, we did not find a significant effect of the indirect haptic disturbance on motivation in this study.

The lack of benefits of the indirect perturbing forces to enhance motor learning may be explained by the extent to which they affect a learner’s internal planning noise and execution noise. Planning noise, the component of motor variability originating in the brain, is thought to be beneficial for motor learning [18]–[20]. The brain has no direct way of assessing information about variability caused by execution noise, which originates in the peripheral nervous system [1], [2], and is therefore not thought to be beneficial for motor learning. For example, Van Der Vliet et al. [19] used a statespace model to determine that planning and execution noise in a visuomotor adaptation task correlated positively and negatively, respectively, with adaptation rate.

This raises the question as to what extent the perturbing forces affected planning noise and execution noise in the current experiment. Task-relevant feedback, e.g., the visual feedback of the pendulum movements, may restrict planning noise as errors in the brain’s internal model for generating movement can be corrected [18]. However, the perturbations directly altered the internal movement of the system (i.e., altering the swing dynamics), which likely influenced the task-relevant feedback, as well as the planning noise. Execution noise contributes more to motor variability than planning noise, so the lack of skill improvement may be the result of the execution noise counteracting the learning benefits from increased planning noise. This relationship was not tested here, but should be explored with similar modeling techniques or careful experimental design.

The perturbations may have also changed the participants’ coordination strategy. The redundancy in the movements may have led participants to adopt a strategy that was either not used in the unperturbed trials or one that was generally less effective. The training period was also relatively short compared to real-life motor learning, so these results may simply reflect incomplete adaptation [6].

### C. Indirect haptic disturbances differentially affects transfer

We found differential effects of the perturbations on the transfer of learning depending on the type of skill learned. Indirect haptic disturbance appears to hamper the transfer to tasks that require a different path to be executed while promoting transfer to a task with altered dynamics. This performance improvement was only present in the short term in the second transfer task, suggesting that indirect haptic disturbance enhances immediate transfer of learning to tasks with altered dynamics.

The benefit of indirect haptic disturbance on transfer to a task with altered dynamics may be explained by participants’ familiarity with unexpected faster movements of the pendulum. Shortening the pendulum rod increased its natural frequency, which might be a similar experience to the perturbations causing a sudden increase of the pendulum angle during training. This is in line with Henry’s Specificity hypothesis, which states that transfer of skill is generally low because different motor tasks require several different motor abilities, which are independent of each other [13]. The transfer task with altered target positions might have required more motor abilities in common with training without indirect haptic disturbances, thereby explaining how participants in the control group better learned to decrease the power at natural frequency in this transfer task.

### D. Indirect haptic disturbance does not affect motivation, but may hamper participants’ sense of agency

According to Wulf and Lewthwaite’s OPTIMAL theory [9], enhanced skill learning can be attributed to an increase in motivation. Perturbations did not appear to considerably alter participants’ motivation. This may encode a ceiling effect, as some scores (e.g., *Interest/Enjoyment*) were already high at baseline.

Sense of agency has also been suggested to be associated with skill learning [11]. The fact that the interaction between *Group* and *Time* was trending towards significance from baseline to training, suggests that the perturbations might have negatively affected participants’ sense of agency, but we cannot currently draw any conclusions on this point. Yet, there was no interaction effect at long-term retention, suggesting that this method could be used in training programs without a lasting effect on participants’ sense of agency.

### E. Future work

The possible relationship between motor variability and planning and execution noise should be studied further. The proposed task, while more ecologically valid due to the incorporation of the system dynamics compared to other lab-based tasks, is likely too complex and may not be suitable for model-fitting methods like those employed by other groups [19]. Simplifying the task, e.g., to only one target, might aid in this regard, although there is no guarantee that these results would be generalizable to real-world motor learning.

Attention is another psychological factor that could influence learning [9]. An external focus of attention is desirable for enhanced performance and skill learning, so future work could include questionnaires to directly quantify this.

## V. Conclusion

Perturbing forces applied to the internal degree of freedom of a dynamic system to be controlled increases motor variability. However, this increased motor variability does not seem to enhance learning of the trained task, compared to training without perturbations. Yet, participants who trained with the indirect haptic perturbations showed better learning transfer in a task that included a modified dynamic system. This came at the cost of worse performance in the transfer task with altered target positions compared to the control group. Importantly, while the haptic disturbance seemed to reduce the participants’ sense of agency, it did not affect the overall participant’s motivation, ruling out motivation as the limiting factor in motor learning experiments with haptic disturbance. This study is the first to use haptic disturbance applied to the internal degree of freedom of the controlled system to increase motor variability, which could be used in future work to increase the exploration of task dynamics.

